# PIE-FLIM measurements of two different FRET-based biosensor activities in the same living cells

**DOI:** 10.1101/2020.01.10.902213

**Authors:** C.A. Reissaus, K.H. Day, R.G. Mirmira, K.W. Dunn, F.M. Pavalko, R.N. Day

## Abstract

We report the use of pulsed interleaved excitation-fluorescence lifetime imaging microscopy (PIE-FLIM) to measure the activities of two different biosensor probes simultaneously in single living cells. Many genetically encoded biosensors rely on the measurement of Förster resonance energy transfer (FRET) to detect changes in biosensor conformation that accompany the targeted cell signaling event. One of the most robust ways of quantifying FRET is to measure changes in the fluorescence lifetime of the donor fluorophore using fluorescence lifetime imaging microscopy (FLIM). The study of complex signaling networks in living cells demands the ability to track more than one of these cellular events at the same time. Here, we demonstrate how PIE-FLIM can separate and quantify the signals from different FRET-based biosensors to simultaneously measure changes in the activity of two cell signaling pathways in the same living cells in tissues. The imaging system described here uses selectable laser wavelengths and synchronized detection gating that can be tailored and optimized for each FRET pair. Proof-of-principle studies showing simultaneous measurement of cytosolic calcium and protein kinase A activity are shown, but the PIE-FLIM approach is broadly applicable to other signaling pathways.

**STATEMENT OF SIGNIFICANCE:** Here, we demonstrate that PIE-FLIM can separate and quantify the signals from two different FRET-based biosensors expressed in the same cells in intact tissues. PIE imaging excites the sample with two pulsed lasers of different wavelengths. The individual excitation pulses are delayed relative to one-another so that they are interleaved at the sample, and the detection channels are synchronized to the laser pulses to permit the discrete measurement of two different probe lifetimes. This enables the independent quantification of changing signals from two FRET-based biosensors. The advantage of PIE-FLIM for multiplexed imaging of FRET-based biosensor probes is that the different donor emission signals are separated in time as well as in spectral space minimizing the problem of crosstalk.

## INTRODUCTION

Cells exist in a dynamic equilibrium with their environment and respond to external perturbations through networks of intracellular signaling pathways. These networks of pathways are interconnected and integrated, allowing them to coordinate signal transduction. The ability to monitor the activities of different cell signaling pathways inside living cells became possible with the development of the genetically encoded biosensor probes (1–4). A variety of design strategies have been used in the development these genetically encoded probes, and many rely on the measurement of Förster resonance energy transfer (FRET) to detect the changes in biosensor conformation that accompany the targeted signaling event (4).

The genetically encoded FRET-based biosensor probes have a modular design consisting of sensing and reporter domains. Typically, the reporter domain consists of a pair of fluorescent proteins (FPs) that share significant spectral overlap that is necessary for efficient FRET. The sensing domain serves as a linker between the FPs and includes an element that is modified by the targeted biological event, as well as a binding motif that recognizes that modification. This allows the sensing domain to change its conformation in response to a specific cell-signaling event, altering the distance between the FP pair in the reporter (1–4). The changing intramolecular FRET signal from these single chain biosensor proteins can be monitored in real time, allowing measurement of the spatiotemporal dynamics of signaling events inside living cells.

Given that networks of cell-signaling pathways are highly integrated, it has long been a goal to develop approaches that allow detection of more than one of these cellular events at the same time (multiplexing, ref. 5,6). Because FRET-based biosensors rely on sensing units that contain two FPs, multiplexing more than one biosensor in the same living cells is difficult because of the limited spectral space (4). There are mathematical approaches for separating multiple fluorescence signals, but these can be challenging when applied to sensitized emission measurements of FRET (7). An alternative approach is to use fluorescence lifetime microscopy (FLIM) to detect FRET (8). Whereas sensitized emission measurements require mathematical corrections to determine FRET, FLIM directly quantifies the decrease in the donor’s fluorescence lifetime that results from energy transfer (7–11). Thus, FLIM-based measurements of two different FRET biosensors require only separation of the two donor signals. In this study, we demonstrate how pulsed interleaved excitation (PIE), when combined with FLIM, can simultaneously measure FRET from two different biosensors in the same living cells.

PIE imaging excites the sample with two pulsed lasers of different wavelengths that are synchronized on the nanosecond scale (12). The pulses from the two different lasers are delayed relative to one-another such that the individual pulses are interleaved at the sample. The detection channels are synchronized to the laser pulses to permit the discrete measurement of two different lifetimes at the same time (12–14). The advantage of PIE-FLIM for multiplexed imaging of FRET-based biosensor probes is that the different donor emission signals are separated in time as well as in spectral space minimizing the problem of crosstalk.

The rationale for developing the PIE-FLIM approach is to enable simultaneous monitoring of two different signaling pathways in living cells and tissues. For example, insulin secretion by pancreatic α-cells is regulated by the coordinated activity of canonical signaling pathways mediated by both intracellular 3’,5’-cyclic adenosine monophosphate (cAMP) and intracellular calcium (Ca^2+^) (15). Because of the central importance of these two signal transduction pathways, much effort has been invested in the development and refinement of biosensors to monitor their activities (4). For example, the "A kinase Activity Reporter" (AKAR) is a single chain FRET-based biosensor for protein kinase A (PKA) signaling in response to cAMP that has undergone significant refinement over the years (1–3). Similarly, the genetically encoded FRET-based biosensors for calcium have also been extensively engineered, with the ‘Twitch’ sensors containing troponin C (TnC) calcium binding domains being among the most sensitive reporters of Ca^2+^ signaling (16). Here, we demonstrate that PIE-FLIM can resolve the signals from the AKAR4 and Twitch2b FRET biosensors in the same cells and provide proof-of-principle studies for the simultaneous measurement of PKA and Ca^2+^ activities in intact pancreatic islets.

## MATERIALS AND METHODS

### FP purification

The purified FPs were produced from plasmids encoding the proteins tagged with the (6) His epitope (provided by Michael Davidson, FSU, Tallahassee, FL). The (6) His-tagged proteins were grown in bacteria, which were then disrupted by sonication, and the supernatant was recovered by centrifugation. The supernatant was mixed with Talon resin, and the eluted proteins were concentrated using an Amicon Ultra centrifugal filter (MilliporeSigma, St. Louis, MO) in PBS (pH 7.4). Aliquots of the concentrated FPs in solution were added to an eight-chambered coverglass for FLIM measurements.

### FRET Standards and Biosensors

The FRET standards were produced using either monomeric (m)Turquoise2 (17) or mNeonGreen (18) as the donor for mScarlet (19). These donor fluorophores were coupled directly to mScarlet through a 10 amino acid (aa) linker as described earlier (20). The FRET standards were used to verify the PIE gating settings for the biosensor measurements. The standards also served as the starting point for the generation of the modified AKAR sensor of PKA activity, and the modified Twitch calcium sensor.

The sensing units of AKAR4 (3) and Twitch2B (16) were obtained by polymerase chain reaction (PCR) and the products were used to replace the linker sequence in the FRET standards. All plasmid inserts were confirmed by direct sequencing.

### Cell culture and transfections

MLO-Y4 osteocytes (21) were cultured on collagen-coated plates (rat tail collagen type I, BD Biosciences, San Jose, CA) in MEM-a (Gibco, Life Technologies, Carlsbad, CA) supplemented with 5% FCS, 5% fetal bovine serum (FBS) and 1 % penicillin/streptomycin (Gibco, Life Technologies,

Grand Island, NY). The cells were harvested at 70-80% confluence and transfected by electroporation at 250 volts with a 9 ms pulse duration using a BTX ECM 830 electroporator (Harvard Apparatus, Holliston, MA) as described earlier (22). The cells are immediately recovered from the cuvette and diluted in phenol red-free tissue culture medium containing serum. The suspension is transferred to sterile two or four well-chambered coverglass (Lab-Tek II, Thermo Scientific, Waltham, MA), which are placed in an incubator (37°C and 5% CO2-Air 95%) prior to FLIM the following day.

### Islet preparation and viral infections

All mouse experiments were performed with approval and oversight from the Indiana University Institutional Animal Care and Use Committee (IACUC). All experiments were performed in accordance with relevant guidelines and regulations. Male C57BL/6J mice 8 weeks of age (Jackson Labs, Bar Harbor, ME) were utilized. To isolate islets, mice were euthanized, the pancreas was harvested, and islets were liberated using a 0.3% collagenase digestion in 37 °C water bath in Hanks buffered sodium salt. Islets were maintained in islet media containing phenol free RPMI 1640, with 10% FBS, 100 U/ml Penicillin, 100 μg/ml Streptomycin, and 8 mM glucose. Islets were allowed to recover overnight prior to adenoviral infection. To infect β-cells *in vitro,* isolated islets were washed with Dulbecco’s phosphate buffered saline without calcium or magnesium, followed by distention with Accutase (MilliporeSigma, St. Louis, MO) at 37 °C for 30 seconds. Accutase was rapidly inactivated with room temperature islet media, then washed with fresh islet media. Adenovirus was added directly to islet media to achieve approximately 1 × 10^8^ viral particles per 150 islets in 1 mL of islet media. Viral infection lasted at least 6 hours. After viral infection, islets were plated in Ibidi μ-Slide 8 well glass bottom slides with fresh islet media. On the day of imaging (24 hours after infection), islets were equilibrated in 2.8mM glucose phenol-free Krebs-Ringer bicarbonate buffer with 10 mM HEPES (KRBH) for 30 minutes. Baseline images were collected, followed by treatment with 1 μM thapsigargin or 2.5 μM forskolin treatments in 2.8mM glucose KRBH. Images were collected at 15 minutes post treatment. For each islet, a z-stack 32 μm thick, with step intervals of 8 μm was collected to sample 4 different planes of cells.

### PIE-FLIM measurements

The fluorescence lifetime measurements are made using the ISS Alba 5 FastFLIM system (ISS Inc., Champagne, IL) coupled to an Olympus IX71 microscope equipped with a 60 X, 1.2 numerical aperture water-immersion objective lens. A Pathology Devices (Pathology Devices, San Diego, CA) stage-top environmental control system maintains the temperature at 37°C and 5% CO_2_-95% Air. The two-channel Alba laser scanning confocal system is controlled by ISS VistaVision software, which allows independent adjustment of the pinholes, and positioning of the detectors for both channels. The system is equipped with a 5 mW 448 nm diode laser and a 200 mW Supercontinuum white light laser (WLL). For the WLL, wavelengths below 690 nm are passed to a filter wheel with excitation filters for green (510/10 nm), or red (561/14 nm). The system uses a 445/515/561 triple-notch filter (Semrock, Rochester, NY) for dual laser excitation.

For PIE-FLIM the 448 nm diode laser and WLL are both modulated at a fundamental frequency of 20 MHz, with additional measurements at up to 8 sinusoidal harmonics (20-180 MHz). Both lasers are combined in the laser launcher and delivered by a single-mode polarization-maintained optical fiber to the confocal scanning system, which is controlled by the VistaVision software (4.2, Build 163, ISS Inc., Champagne, IL). The firmware allows independent adjustment of channel gating and pulse width for PIE-FLIM. The fluorescence signals emitted from the specimen are routed by a 495 nm long pass beam splitter through 474/23 nm (cyan emission) or 543/22 nm (green emission) band-pass emission filters, and the signals are detected using two identical avalanche photodiodes (APD). The software records the time resolved photons in different cross frequency phase bins to generate the phase histogram. The information at higher harmonics is obtained by the digital frequency transform of the phase histogram. The phase delays and modulation ratios of the emission signal are measured at each pixel of an image for each frequency. The cyan channel is calibrated with 5μM solution of Atto 425 NHS ester (MilliporeSigma, St. Louis, MO) dissolved in water (lifetime 3.6 ns, ref. 23) and the green channel is calibrated with 5 μM solution of fluorescein (MilliporeSigma, St. Louis, MO) dissolved in 1 mM NaOH (4 ns lifetime, ref. 24). The PIE gating settings were then verified using a 50:50 mixture of the two dyes.

## RESULTS

### PIE-FLIM

The basic concept of PIE-FLIM imaging is illustrated in **Fig. 1**. The PIE-FLIM system used here modulates both a 448 nm and a white light laser (WLL) at the same rate of 20 MHz, resulting in a pulse repetition period of 50 ns. The pulse trains from each laser are temporally offset from one-another, permitting the individual pulses to be interleaved when arriving at the sample (**Fig. 1** *A*). The APD detectors for the two emission channels are synchronized with the laser clock, allowing measurement of the fluorescence decay for two different fluorophores immediately following the corresponding excitation laser pulse. The interleaved illumination and detection scheme enable the separation of two different fluorescent signals into discrete gates (**Fig. 1** *B*).

**Figure 1.**
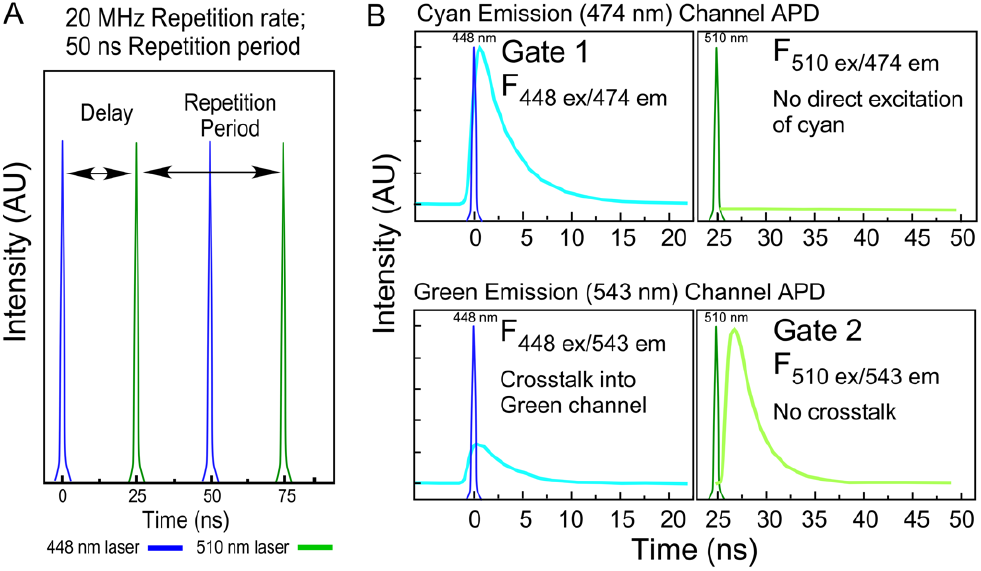
Schematic representation of PIE-FLIM. (*A*) This study uses both a 448 nm diode laser (blue pulse) and a white light laser filtered to 510 nm (green pulse). The repetition rate for both lasers is 20 MHz, resulting in a 50 ns repetition period. The pulse trains are delayed relative to one-another to interleave the pulses at 25 ns intervals. *(B)* The detection channels are synchronized to the laser clock, such that the emission decay produced by each excitation immediately follows the laser pulse. The gating windows (gate 1, gate 2) are optimized for the collection of photons in each channel. Note that cyan crosstalk into the green channel is not detected in gate 2 because the 448 nm excitation is separated in time from the detection of the green emission.

The laser excitation, filter selection and PIE gating described here are designed to separate the fluorescence signals from the genetically encoded mTurquoise2 (17) and mNeonGreen (18) FPs. The accurate measurement of lifetime decays requires an interval of at least 5 lifetimes (5*τ*) before the next excitation pulse (25). The imaging system software (VistaVision, ISS) enables the independent adjustment of the delay and pulse width to optimize photon collection efficiency in PIE gate 1 for mTurquoise2 (448 nm excitation and 474/23 nm emission) and PIE gate 2 for mNeonGreen (510 nm excitation and 543/22 nm emission). The delay between the laser pulses is set to 25 ns, which is a sufficient interval to measure the nearly complete decay of the FPs used in this study (**Fig. 1** *B*). Importantly, mTurquoise2 emission crosstalk into the mNeonGreen detection channel (gate 2) is avoided, since the two emission signals are separated in time from one-another (**Fig. 1** *B*).

### Phasor Plot Analysis of Lifetime Distributions

The accurate measurement of lifetimes by frequency domain FLIM requires calibration with standards that have spectral characteristics and lifetimes that are similar to the FPs used for the experiments. Here, Atto 425 (Ex 439, Em 485, lifetime 3.6 ns, ref. 23) is used as the standard for gate 1 and fluorescein (Ex 490, Em 514, lifetime 4 ns, ref. 24) is used as the standard for gate 2. The phase histogram of the fluorescence response for Atto 425 excited with the 448 nm laser shows the exponential decay of fluorescence that is detected in gate 1 (Em 474/23 nm). The phasor plot analysis for Atto 425 demonstrates a single exponential decay with a lifetime distribution centered at 3.6 ns (Supplemental **Fig. S1** *A*). Similarly, the phase histogram for the fluorescence response of fluorescein excited at 510 nm shows the fluorescence decay that is detected in gate 2 (Em 543/22 nm), and the phasor plot indicates an average single-component lifetime of 4 ns (Supplemental **Fig. S1** *B*). PIE-FLIM is then used to measure the lifetimes of both Atto 425 and fluorescein in a 50:50 mixture of the dyes. The dye mixture is being illuminated with both lasers, and the detectors for gate 1 and gate 2 are synchronized with the laser clock (see **Fig. 1**). Supplemental **Fig. S1** *C* shows that the phase histograms for the fluorescence decays of both Atto 425 and fluorescein are well separated, and the individual lifetime distributions for the two dyes are clearly resolved on the combined phasor plot.

To verify that the PIE-FLIM gating is optimally set for detection and discrimination of the donor FPs used in this study, simultaneous lifetime measurements were obtained from a mixture of purified (6) His-tagged-mTurquoise2 and -mNeonGreen in phosphate buffered saline. The mixture of the two purified FPs was illuminated with both lasers, and the emissions were detected in the two PIE gates. Supplemental **Fig. S1** *D* shows that the phase histograms of the fluorescence decays for the two purified FPs are well separated by the PIE gating. The combined phasor plot resolves the lifetime distributions for both, yielding the expected lifetimes for the unquenched FPs in solution (Supplemental **Table S1**).

### PIE-FLIM measurements of mTurquoise2 and mNeonGreen in living cells

Next, MLO-Y4 cells, which have characteristics of osteocytes (21), were transfected by electroporation with mammalian expression plasmids encoding either mTurquoise2 or mNeonGreen. Where indicated, cells transfected with either mTurquoise2 or mNeonGreen were mixed immediately after electroporation in the same chambered coverglass, or alternatively co-transfected with the two different plasmids. **Fig. 2** *A* shows PIE-FLIM measurements of a cell expressing only mTurquoise2, which is detected in gate 1. The phasor plot from gate 1 indicates an average single-component lifetime of 3.97 ns. Measurements acquired from multiple cells expressing mTurquoise2 are presented in Supplemental **Table S1**. By contrast, a cell expressing only mNeonGreen is detected in gate 2, and the phasor analysis reveals an average single component lifetime of 3.07 ns (**Fig. 2** *B* and Supplemental **Table S1**). It is known that quenching events in the living cell environment affect the lifetime of the FPs, causing the lifetimes to be shorter than those measured for the purified FPs in solution (26).

**FIGURE 2.**
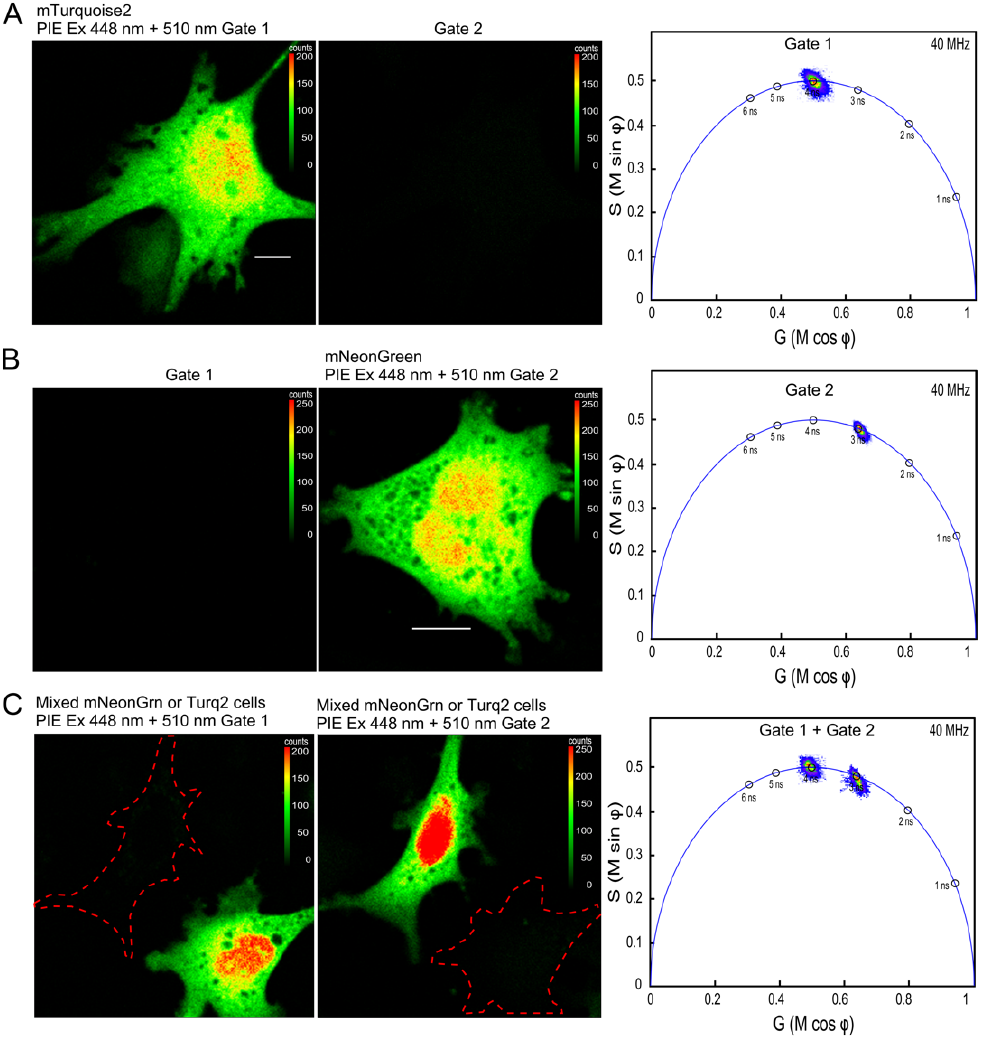
PIE-FLIM of cells expressing either mTurquoise2 or mNeonGreen. (*A*) Cell expressing mTurquoise2 was illuminated with both 448 nm and 510 nm laser lines, and the phasor analysis for gate 1 (Ex 448 nm, Em 474/23 nm) is shown. The lookup table (LUT) indicates intensity in counts per pixel and the scale bar is 10 μm in length. (*B*) PIE-FLIM measurement of a cell expressing mNeonGreen, showing the phasor analysis for gate 2 (Ex 510 nm, Em 543/22 nm). (*C*) PIE-FLIM measurement cells that were transfected with either mTurquoise2 or mNeonGreen and mixed in the chambered coverglass. A FOV with two cells in which one is expressing mTurquoise2 (left panel, gate 1) and the other is expressing mNeonGreen (right panel, gate 2) is shown. The dashed outline indicates the position of the cell in the FOV that is not detected in the gate. The combined phasor plot shows the lifetime distributions measured for each gate.

To demonstrate that PIE-FLIM can measure the lifetimes of both FPs simultaneously, cells expressing each FP were mixed on the same chambered coverglass. A field of view (FOV) with two adjacent cells expressing either mNeonGreen or mTurquoise2 was identified, and PIE-FLIM measurements were acquired. In **Fig. 2** *C*, the cell expressing mTurquoise2 (left panel) is only detected in gate 1, whereas the cell expressing mNeonGreen (right panel) is detected only in gate 2. The phasor plots obtained for each gate clearly resolve the individual lifetimes for either mTurquoise2 or mNeonGreen. The average lifetimes for mNeonGreen and mTurquoise2 are presented in Supplemental **Table S1**.

Next, to demonstrate that PIE-FLIM can quantify the individual lifetime distributions for both mTurquoise2 and mNeonGreen in the same cell, MLO-Y4 cells were co-transfected with plasmids encoding both FPs. A FOV with cells co-expressing the two different FPs was identified, and PIE-FLIM measurements were acquired (**Fig. 3**). All cells are detected in both gates because each coexpresses the two FPs to differing degrees. The phasor plots obtained in each gate clearly resolve the different lifetime contributions from both mNeonGreen and mTurquoise2. The average lifetime for the co-transfected mNeonGreen was 3.09 ns, and average lifetime for mTurquoise2 was 3.89 ns (**Fig. 3, Table S1**). Together, these results show that PIE-FLIM gating can separate the individual lifetime contributions from both mNeonGreen and mTurquoise2 produced in the same living cells.

**FIGURE 3.**
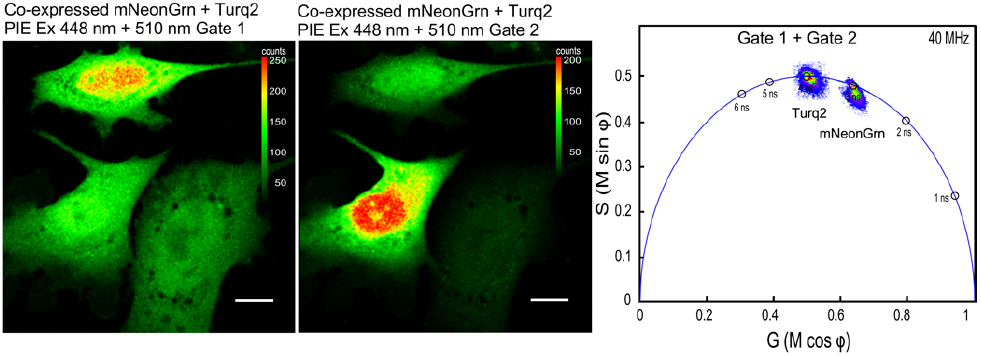
PIE-FLIM of cells co-expressing mTurquoise2 and mNeonGreen. A FOV with cells that all co-express both mTurquoise2 (left panel, gate 1) and mNeonGreen (right panel, gate 2) is shown. The LUT indicates intensity in counts per pixel and the scale bar is 10 μm in length. The combined phasor plot shows the lifetime distributions that are measured for each gate.

### PIE-FLIM measurements of two different FRET standards

The goal of this study is to use the PIE-FLIM to measure the activities of two different FRET-based biosensors that consist of either mTurquoise2 or mNeonGreen as the FRET donors for mScarlet I. Both donor FPs share sufficient spectral overlap with mScarlet for efficient energy transfer (27, 28). To demonstrate the feasibility of this approach studies were conducted using FRET standards (20) consisting of either mTurquoise2 or mNeonGreen coupled directly to mScarlet I through a 10aa linker (FRET 10; see **Fig. 4**). Cells expressing either mTurquoise2-Scarlet FRET 10 or mNeonGreen-Scarlet FRET 10 fusion proteins were imaged using PIE-FLIM, and the FRET efficiency (EFRET) for each of the standards was quantified. Additionally, as described above, cells transfected with plasmids encoding either of the FRET 10 standards were also mixed in the same chambered coverglass.

**FIGURE 4.**
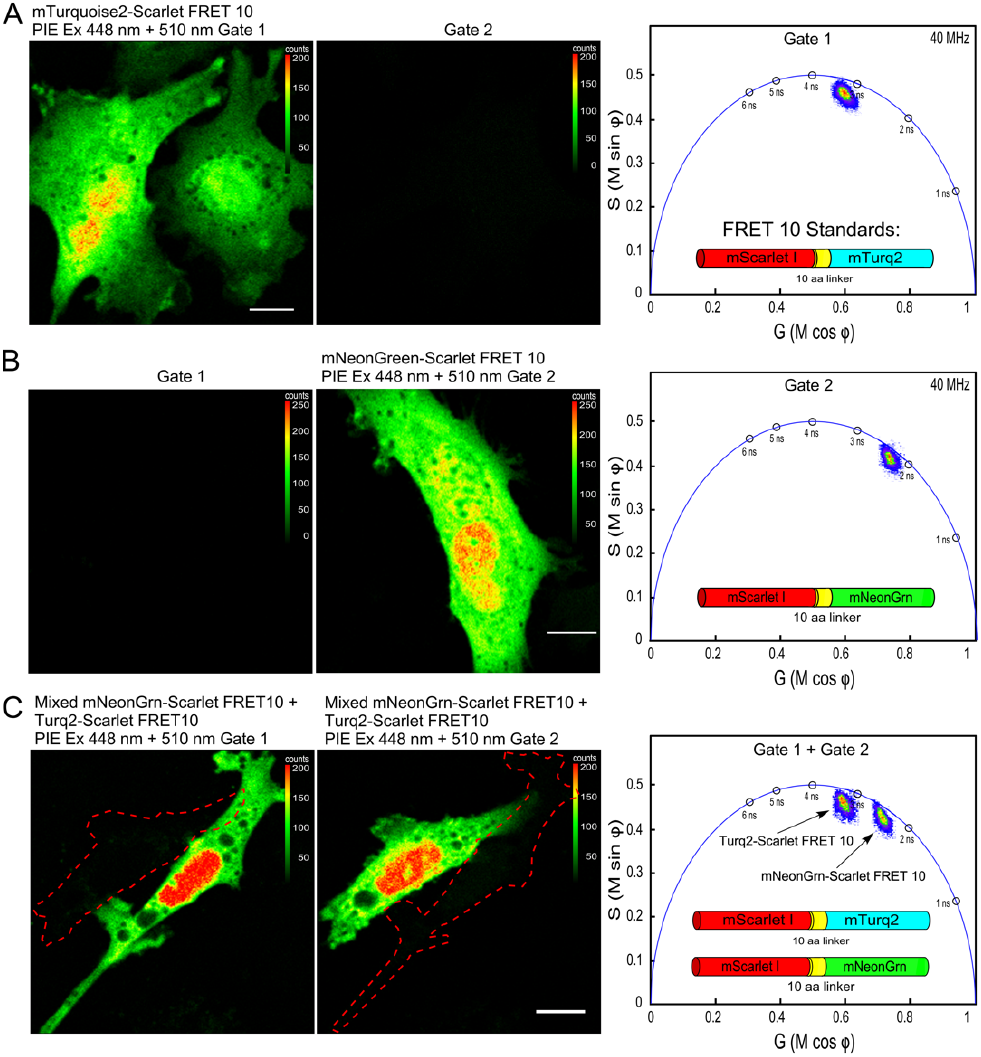
PIE-FLIM of cells expressing either mTurquoise2-Scarlet or mNeonGreen-Scarlet FRET 10 standards. (*A*) Measurements from cells that express mTurquoise2-Scarlet FRET 10. The LUT indicates intensity in counts per pixel and the scale bar is 10 μm in length. The phasor plot shows the lifetime distribution for gate 1. (*B*) Measurement of a cell that expresses mNeonGreen-Scarlet FRET 10. The phasor plot shows the lifetime distribution for gate 2. (*C*) PIE-FLIM measurements from cells that were transfected with either of the FRET standards and mixed in the chambered coverglass. A FOV with two cells, one expressing mTurq2-Scarlet FRET 10 (left panel, gate 1) and the other expressing mNeonGrn-Scarlet FRET 10 (right panel, gate 2) is shown. The outline of the cell not detected in the respective gate is indicated. The combined phasor plot shows the lifetime distributions measured for each gate.

In **Fig. 4** *A*, cells expressing the mTurquoise2-Scarlet FRET 10 fusion protein were imaged using PIE-FLIM and detected in gate 1. The phasor plot shows that the lifetime is quenched compared to the lifetime for mTurquoise2 alone (compare with **Fig. 2** *A* and Supplemental **Table S1**), and the lifetime distribution is shifted inside the universal semicircle, indicating a multi-exponential decay consistent with FRET (22). The average lifetime is 2.5 ns, corresponding to a FRET efficiency of 37% (Supplemental **Table S2**). Similarly, the phasor analysis for cells expressing the mNeonGreen-Scarlet FRET 10 standard detected in gate 2 also indicates a quenched, multi-exponential decay consistent with FRET (**Fig. 4** *B*; compare with **Fig. 2** *B*). The average lifetime is 2.1 ns, corresponding to a FRET efficiency of 33% (Supplemental **Table S2**).

To demonstrate that the signals from the two different FRET standards can be separated by PIE gating, cells that were mixed after transfection with either of the two FRET 10 fusion proteins were imaged. A FOV with adjacent cells expressing either the mTurquoise2-Scarlet FRET 10 or the mNeonGreen-Scarlet FRET 10 fusion proteins was identified, and PIE-FLIM measurements were acquired. The cell expressing mTurquoise2-Scarlet FRET 10 (left panel) and the cell expressing mNeonGreen-Scarlet FRET 10 (right panel) are detected in their respective gates (**Fig. 4** *C*). The phasor plots obtained for each gate clearly resolve the individual lifetimes for two different FRET standards (**Fig. 4** *C* and Supplemental **Table S2**).

Next, MLO-Y4 cells co-expressing the two different FRET standards were imaged using PIE-FLIM. Here, individual cells that co-express both FRET 10 fusion proteins are detected in both gates because (**Fig. 5**). The phasor plots obtained for each gate clearly resolve the individual quenched lifetimes for both mTurquoise2-Scarlet FRET 10 and mNeonGreen-Scarlet FRET 10 in each individual cell (**Fig. 5** and Supplemental **Table S2**). Taken together, the results demonstrate that PIE-FLIM can simultaneously quantify the quenched lifetimes for two different FRET probes inside the same living cells.

**FIGURE 5.**
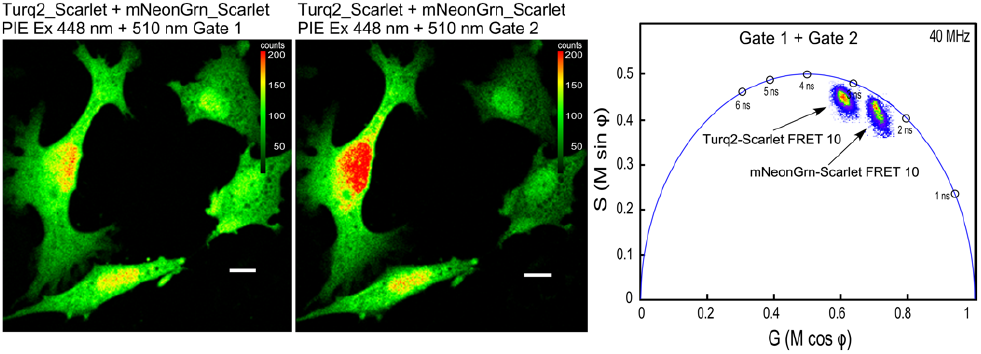
PIE-FLIM of cells co-expressing both mTurq2-Scarlet and mNeonGrn-Scarlet FRET 10 standards. A FOV with cells that co-express both mTurq2-Scarlet FRET 10 (left panel, gate 1) and mNeonGrn-Scarlet FRET 10 (right panel, gate 2) is shown. The LUT indicates intensity in counts per pixel and the scale bar is 10 μm in length. The combined phasor plot shows the lifetime distributions that are measured for each gate.

### PIE-FLIM measurements of two different biosensors in the cells of intact pancreatic islets

The FRET 10 standards were packaged into Adenovirus and transduced into isolated mouse pancreatic islets. The production of the FRET standards is under the control of the viral CMV (cytomegalo virus) promoter, so they are expected to be expressed in all transduced islet cells. The cells within the intact islets that expressed the mTurquoise2-Scarlet FRET 10 standard had an average quenched lifetime of 2.81 ns, corresponding to a FRET efficiency of 28%. Similarly, the cells expressing the mNeon-Scarlet FRET 10 standard had an average quenched lifetime 1.97 ns, corresponding to a FRET efficiency of 34% (Supplemental **Table S3**). The analysis of cells in the intact islets that co-expressed of both standards showed the same distribution of quenched lifetimes as that of either standard alone (**Table S3**).

Since both cAMP and calcium pathways are important regulators of insulin secretion from β-cells in the pancreatic islets (15), the PKA biosensor AKAR (1–3) and the Ca^2+^ biosensor Twitch 2b (16) were modified to incorporate either mTurquoise2 or mNeonGreen as donors, and mScarlet I as the common acceptor. The coding sequences for the modified biosensors were inserted into insulin-promoter driven plasmids (29) for expression specifically in pancreatic α-cells, and used to generate Adenoviral vectors. These were used to transduce the two different biosensors into primary, isolated islets (**Fig. 6**). The intact islets were imaged using PIE-FLIM, and α-cells expressing the mNeonGreen Twitch 2b probe were identified in gate 1 (**Fig. 6** *A*), while α-cells expressing the mTurquoise2 AKAR probe were identified in gate 2 (**Fig. 6** *B*). The α-cells that co-express both probes were identified in both channels (**Fig. 6** *C*), and images were acquired in several different Z-planes, allowing many cells to be analyzed in each islet (**Fig. 6** *D*).

**FIGURE 6.**
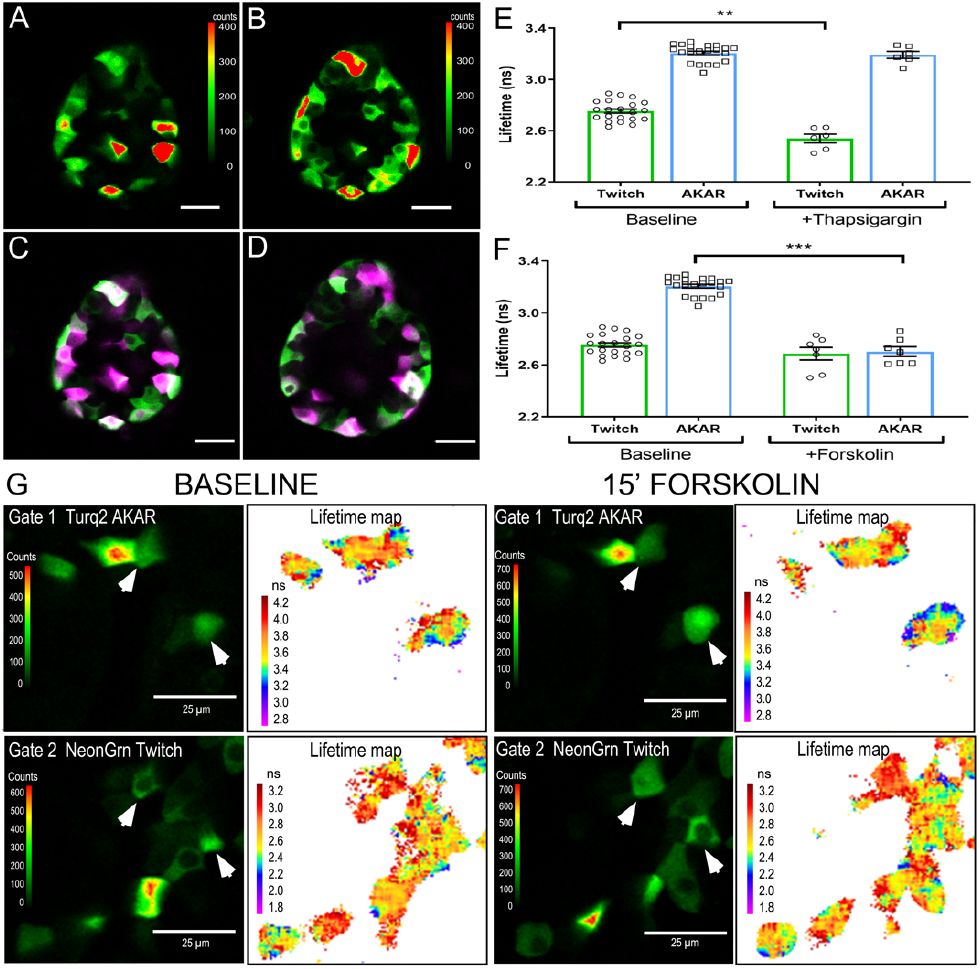
PIE-FLIM of pancreatic islet β-cells expressing both mNeonGrn-Scarlet Twitch2b and mTurq2-Scarlet AKAR4.2 biosensors. An optical section through a representative islet expressing both (*A*) mTurq2-Scarlet AKAR4.2 and (*B*) mNeonGrn-Scarlet Twitch2b biosensors in β-cells. The LUT indicates intensity in counts per pixel and the scale bar is 25 μm in length. (*C*) A pseudo-color merge of panels (*A*) and (*B*) representing AKAR4.2 as magenta and Twitch2b as green. (*D*) The same islet at an optical plane 8 μm above the original plane. Islets co-expressing the both biosensors were treated with (*E*) 1 μM Thapsigargin or (*F*) 2.5 μM forskolin. Each data point represents a single islet with >20 virally transduced cells. (*G*) Changes in biosensor lifetimes were monitored in a different islet transduced with viral vectors for both Turq2 AKAR (top panel) and NeonGrn Twitch2b (bottom panel). After baseline imaging (left), the islet was treated with forskolin for 15 min (right), and images were re-acquired in the same focal plane. α-cells cotransduced with both biosensors are indicated with arrow heads.

To monitor changes in the activity of the two different biosensors the islets were treated with 1 μM thapsigargin or 2.5 μM forskolin to modulate intracellular calcium and cAMP levels, respectively (**Fig. 6** *E and F*). With the addition of thapsigargin, we observed a decrease in the average lifetime for the mNeonGreen Twitch2b probe from 2.75 ns at baseline to 2.53 ns after treatment (**Fig. 6** *E*). There was no measurable change in the mTurquoise2 lifetime within the AKAR4 probe with thapsigargin treatment. With the addition of forskolin, we observed a decrease in the average lifetime for the mTurquoise2 AKAR4 probe from 3.20 ns at baseline to 2.70 ns after treatment (**Fig. 6** *F*). There was no measurable change in the mNeonGreen Twitch 2b lifetime with forskolin treatment (**Fig. 6** *F*). Lifetime maps of α-cells from the same Z-plane of an islet before and 15 min after treatment with forskolin illustrates the change in the AKAR probe lifetime in individual cells in response to the treatment (**Fig. 6** *G*). In contrast, the lifetime map indicates no change in the lifetime of the Twitch 2b probe after forskolin treatment (**Fig. 6** *G*). Together, these results demonstrate PIE-FLIM can separate and quantify the signals from two different biosensors expressed in the same cells in intact tissues.

## DISCUSSION

Here, we describe proof-of-principle studies demonstrating PIE-FLIM measurements of two different biosensor probes in individual cells of intact, living tissue. The use of 20 MHz pulsed lasers permitted a 25 ns delay between PIE pulses, which is a sufficient interval for nearly complete lifetime decay of the FPs used in this study. This temporal gating mitigates the problem of photons arising from one probe affecting the measurements in the gate for the other probe (**Fig. S1** *C and D*). In this regard, the PIE-FLIM measurements of probe activities are best described as "quasi-simultaneous" since the two measurements are separated in time from one-another (12–14). It would be challenging to use higher laser frequencies for this approach, since the repetition period would become shorter than the 5*τ* interval necessary for the measurement of the complete lifetime decay for most FPs (25). Importantly, the PIE-FLIM approach can minimize fluorescence crosstalk between the detection channels because measurements of the emission signals are separated in time from one-another.

The imaging system described here was designed to detect and separate the signals from biosensors based on mTurquoise2 and mNeonGreen, but the combination of the WLL, diode laser, and filters can easily be modified for measurements of other biosensor combinations. We found that the Turquoise2-Scarlet AKAR4 probe used here has a larger dynamic range compared to mNeonGreen-Scarlet Twitch2b in response to drug treatments. This may reflect a limitation of mNeonGreen as a FRET donor; a shortcoming that might be addressed by replacement with a different yellow-green FP. Future permutations of the biosensors could also utilize other fluorescent proteins, including the dark chromoprotein ShadowRed (30), to enhance the FRET efficiency of the probes and to free up spectral space for the addition of other probes.

Of the many imaging methods available to monitor the changing FRET signal from biosensor probes, FLIM is the most quantitative approach (8). However, one shortcoming is that accumulation of sufficient photons for accurate FLIM measurements can be time-consuming. The accurate measurement of probe lifetimes from cells in intact islets required about 10 seconds for the two-channel 256×256 pixel images from a single Z-plane. This is potentially limiting for the detection of fast cellular events such as calcium transients, which are of shorter duration than our acquisition period.

An advantage of using PIE-FLIM measurements to monitor the activities of two different cell signaling events is that both can be independently modulated and quantified. This will be critical to future studies in which novel pharmacological or genetic manipulation might alter one or more regulatory pathways simultaneously. While the levels of intracellular, cytosolic cAMP and calcium act as central regulators of insulin secretion, other pathways are pertinent to β-cell biology. For example, the regulation of calcium levels within the endoplasmic reticulum (ER) is required for insulin production, maturation, and secretion (31). It should be possible to utilize the cytosolic-targeted Twitch2b (kd = 200 nM) and an ER-targeted Twitch2b (54S+, kd = 174 μM) biosensors (16) with the different donor fluorophores to uncover compartment specific differences in calcium regulation. The applications of PIE-FLIM are wide-ranging and increasing as more pathway- and organelle-specific biosensors are developed.

## SUPPORTING MATERIAL

Supporting Material can be found online at

## AUTHOR CONTRIBUTIONS

R.N.D. and C.A.R. conceived and designed the experiments. R.N.D., C.A.R. and K.H.D. carried out the experiments, data analysis, and preparation of the reagents and materials. R.N.D. and C.A.R. wrote the paper. All coauthors discussed the results and exchanged comments on the manuscript.

## ACKNOWLEDGMENTS

The authors thank Drs. Yuansheng Sun and Shih-Chu Liao at ISS, Inc. (Champagne, IL) for their help with the PIE-FLIM hardware and software. This research is supported by the National Institutes of Health RO1 AR052682 (F.M.P), R01 DK060581 and R01 DK105588 (to R.G.M.), the Indiana University School of Medicine (R.N.D), and T32 KD0604466 (C.A.R). This study utilized Diabetes Center core resources supported by National Institutes of Health grant P30 DK097512 (to Indiana University).

